# Leveraging Machine Learning to Uncover the Hidden Links between Trusting Behavior and Biological Markers

**DOI:** 10.1101/2023.09.12.557384

**Authors:** Zimu Cao, Daiki Setoyama, Daudelin Monica-Natsumi, Toshio Matsushima, Yuichiro Yada, Motoki Watabe, Takatoshi Hikida, Takahiro A Kato, Honda Naoki

## Abstract

Understanding the decision-making mechanisms underlying trust is essential, particularly for patients with mental disorders who experience difficulties in developing trust. We aimed to explore biomarkers associated with trust-based decision-making by quantitative analysis. However, quantification of decision-making properties is difficult because it cannot be directly observed. Here, we developed a machine learning method based on Bayesian hierarchical model to quantitatively decode the decision-making properties from behavioral data of a trust game. By applying the method to data of patients with MDD and healthy controls, we estimated model parameters regulating trusting decision-making. The estimated model was able to predict behaviors of each participant. Although there is no difference of the estimated parameters between MDD and healthy controls, several biomarkers were associated with the decision-making properties in trusting behavior. Our findings provide valuable insights into the trusting decision-making, offering a basis for developing targeted interventions to improve their social functioning and overall well-being.

## Introduction

Humans engage in trusting behaviors that rely on their first impressions of others. For example, voting and cooperative activities such as business can be seen as trusting behaviors ^1,2^. However, patients with major mental disorders (MDD) and hikikomori face difficulties in developing trust, which can be interpreted as a defect in decision making^3–5^. Many studies have only described statistical differences in trusting behaviors between patients with MDD and healthy controls (HCs) using standard psychology methods such as behavioral tasks and questionnaires ^6,7^. Understanding decision-making mechanisms that depend on trust, which is latent and not directly observed, is important. However, to the best of our knowledge, no method has been established to decode the degree of trust in decision-making behaviors. Therefore, we developed a machine learning method to model trusting behaviors based on the first impression of others’ facial pictures using a trust game.

A trust game is an economic game used to assess the degree to which people trust others ^8^. The trust game consists of two players, one of whom is asked to evaluate the level of trust in the other and then make a financial decision with a risk. The advantage of using the trust game is that it objectively obtains patients’ interactions, that is, the amount of money the patients pay, in contrast to the traditional diagnosis of psychiatric disorders, most of which are based on subjective clinical interviews. Several clinical studies using economic games have revealed that patients with major depression and personality disorders experience difficulties in social decision-making involving trust ^9,10^. In our previous study using the trust game, we found that male patients with MDD exhibited more trusting behavior toward females of higher attractiveness ^3^; however, most of these studies only conducted descriptive statistical analyses of trust behaviors and did not explore the mechanisms underlying decision making during trust behaviors.

Meanwhile, some blood biomarkers have been shown to change in patients with psychiatric disorders. We previously investigated blood biomarkers in hikikomori patients who avoided social interaction to an extreme extent due to their inability to build a trusting relationship with other people^11^. We found that these patients had decreased blood uric acid levels ^12^. In addition, changes in blood biomarkers have been reported in patients with MDD: a decrease in brain-derived neurotrophic factor (BDNF) and an increase in C-reactive protein (CRP) levels ^13,14^. Furthermore, we demonstrated that minocycline modulates people’s social behavior, suggesting that blood chemical levels play an important in trust behavior ^15^. However, the quantitative relationship between biomarkers and trust behavior remains elusive.

To explore the biomarkers associated with trust behavior, we proposed three biomarker levels. The first level of biomarkers is used to predict latent states, such as healthy conditions or depression, the second is associated with the direct observation of behavioral experiments, and the third reflects the decision-making process (DMP) for each individual. Biomarkers at the first and second levels are easy to determine because latent states and behavioral data are observable. However, the DMP of trusting behavior cannot be directly observed, which can be defined by transformation from the environmental information to behaviors. Therefore, studies on the third level biomarkers related to the DMP are scarce. This study aimed to identify third-level DMP biomarkers. The strategy is as follows: First, we developed a machine learning method to quantitatively decode the DMP in the trust behaviors of both patients with MDD and HCs. This method was based on a Bayesian hierarchical model that describes the decision-making processes in a trust game. Using the MDD and HC data, we estimated the model parameters, which regulated trust behavior. In addition, we quantified the characteristics of trusting behavior in patients with MDD and identified several biomarkers associated with the properties of the MDP involved in trust behavior.

## Results

### Trust game and blood biomarkers

To investigate the relationship between biomarkers and decision-making processes in trust behaviors in patients with MDD, we performed psychological tasks and collected blood biomarkers. Trusting behavior was examined using a trust game (**Fig. 1A**). The trust game we adopted included two players: participants of interest and photographed partners on the PC screen. All participants were asked to score the trustworthiness and attractiveness of the partner using facial information and then make a financial decision on whether to give them some amount of money (**Fig. 1B**). Before playing the game, the participants were initially given some funds and told that their partner had obtained three times the money they were given and would possibly return a portion to them. In addition, all participants provided a non-fasting venous blood sample from which we measured 85 biomarkers (**Fig. 1C**). For comparison, we performed the experiments on patients with MDD and HCs.

**Fig. 1:**
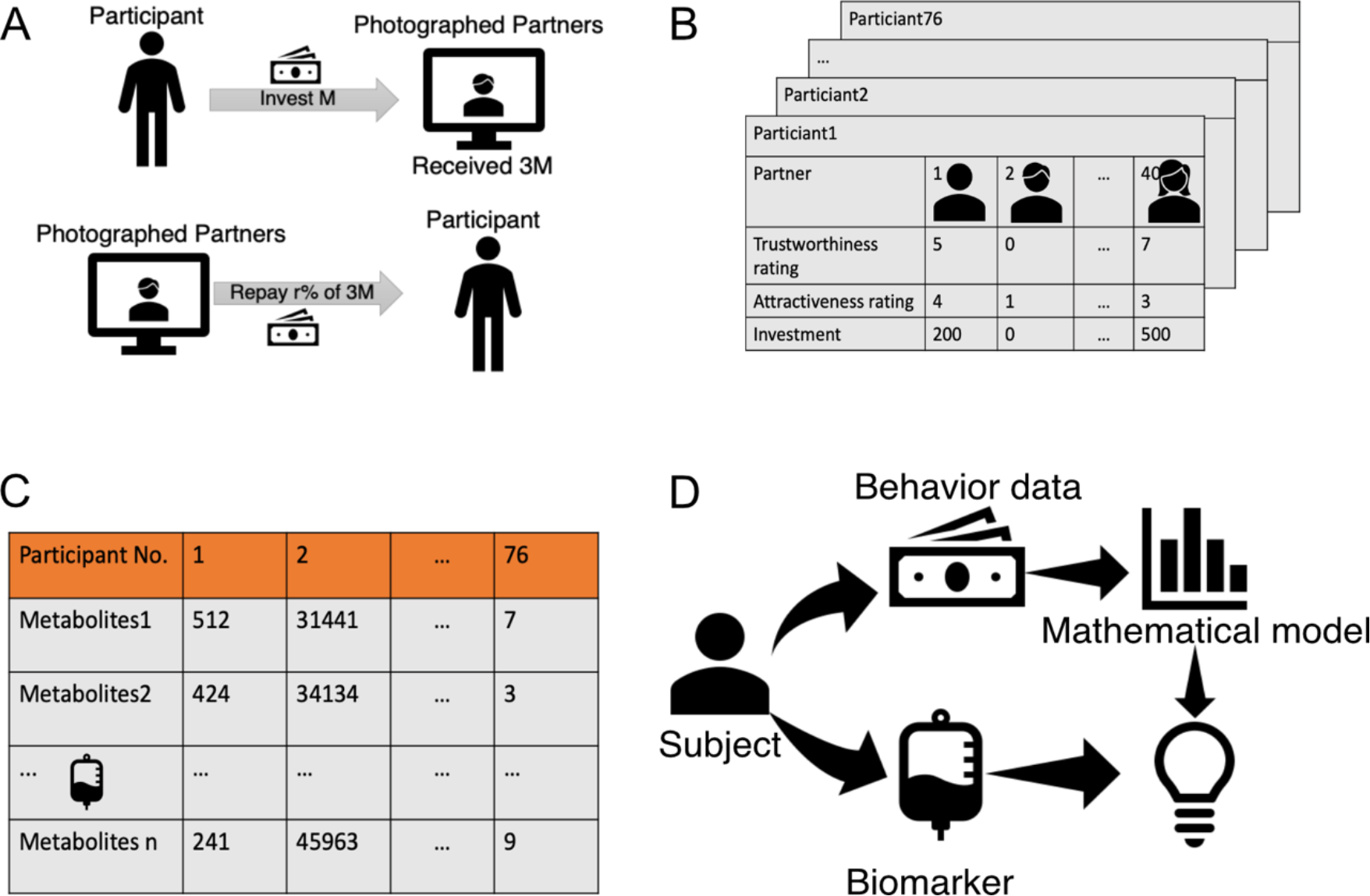
Trust game and blood biomarker measurement. **(A)** Trust game with the photographed partners. A participant is asked to rate the trustworthiness and attractiveness of the partner in the display and to invest 0–1300 yen. **(B)** Behavioral data for the trust game. 38 healthy controls and 38 patients with MDD participated in the trust game. Each participant plays the trust game with 40 photographed partners. **(C)** Measurement of blood biomarkers. Eighty-five metabolites were measured for all participants. The three panels depict examples of metabolite measurement, which show statistically significant differences between HCs and patients with MDD. **(D)** Scheme of this research. For all participants, behavioral data from the trust game and blood biomarker data of 85 metabolites are obtained. From the behavioral data, participant-specific parameters of the decision-making model are estimated. By integrating the estimated parameters and blood biomarker data, we identified the molecular basis of trusting decision making.

### Several types of biomarkers

Many researchers have aimed to identify biomarkers that can predict diseases and healthy states. In general, biomarkers have been considered to predict whether an individual is healthy or in a state of MDD. Previously, we used machine learning to identify biomarkers by classifying HCs and patients with MDD ^11^; patients with MDD had higher concentrations of orotic acid and lactic acid and lower concentrations of cystine, arginine, and methionine in the blood compared to HCs. Another type of biomarkers is indicators that directly predict the participant’s answers and behaviors in the task, which can be identified by a native comparison between behavioral and blood sample data. In fact, we found that participants with higher concentrations of uric acid, carnitine, methionine, creatinine, and nicotinic acid scored higher on the trustworthiness and attractiveness of the photographed partners and invested more money to their partners (**Fig. 2**). However, it remains to be elucidated how these two types of biomarkers are related to the decision-making process in trust behaviors.

**Fig. 2:**
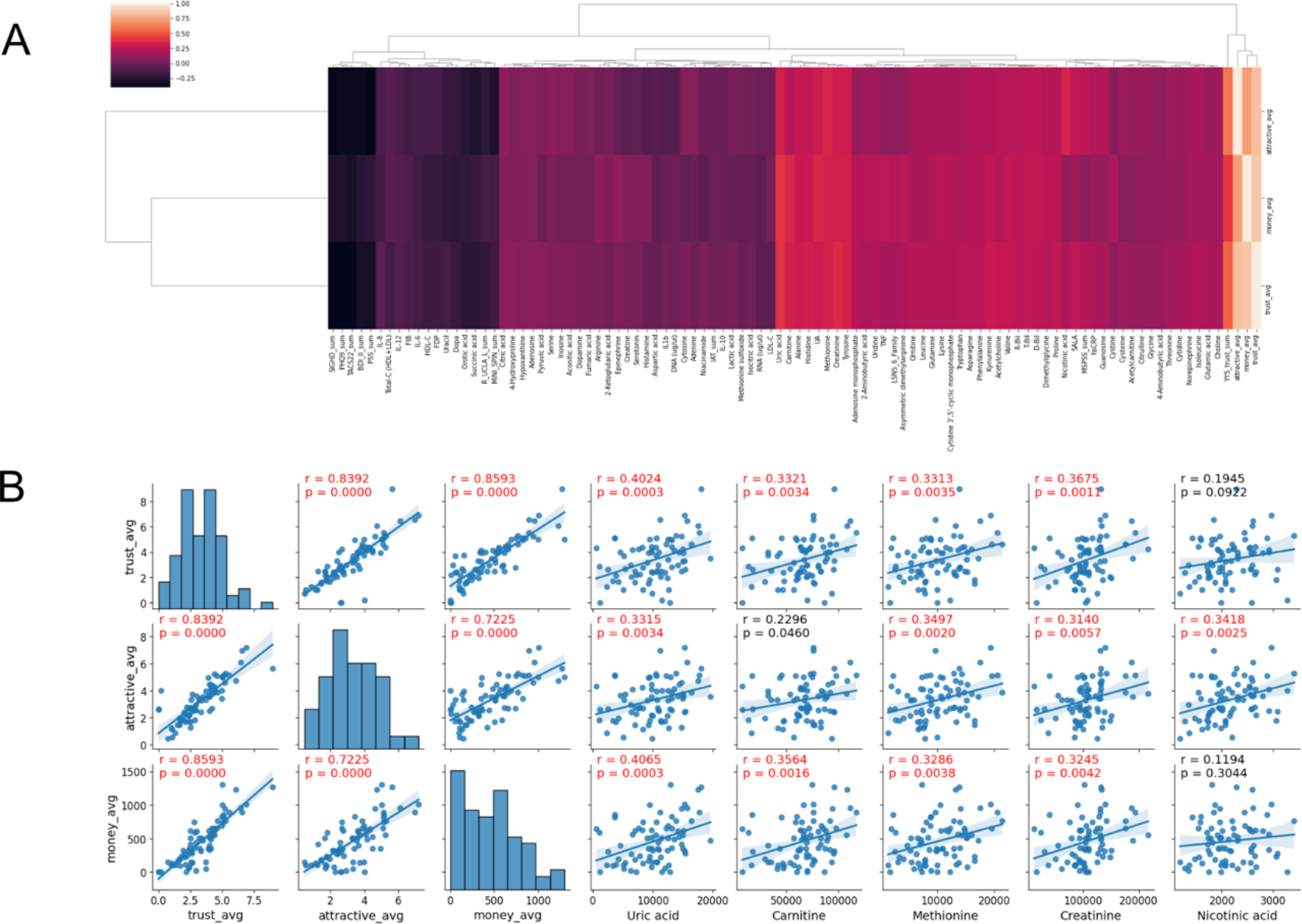
Blood biomarkers correlated with behaviors in the trust game. **(A)** Heatmap of correlation coefficients between the all-estimated parameters and three kinds of behaviors. Hierarchical clustering was performed. Pearson correlation was conducted for correlation analysis. **(B)** Scatter plot of parameters and biomarkers for selected pairs. Correlation coefficients with absolute values > 0.3 and FDR < 0.05 were considered significant, which were colored by red.

In this study, we aimed to identify biomarkers reflecting the decision-making process, that is, the transformation from evaluation of the partner (attractiveness and trustworthiness) to payment behaviors. To this end, we needed to model the decision-making process and estimate the model parameters that reflect the decision-making properties for each participant. The strategy of this study was to integrate data from the trust game and the biomarkers (**Fig. 1D**).

### Bayesian Hierarchical model for trust-based investment

We developed a Bayesian hierarchical model to capture the decision-making process involved in trust-based investments (**Fig. 3A**). Specifically, our model considers the evaluation of a partner’s face to be a crucial determinant of investment decisions. In the model, the participant evaluated the partner *j* as

**Fig. 3:**
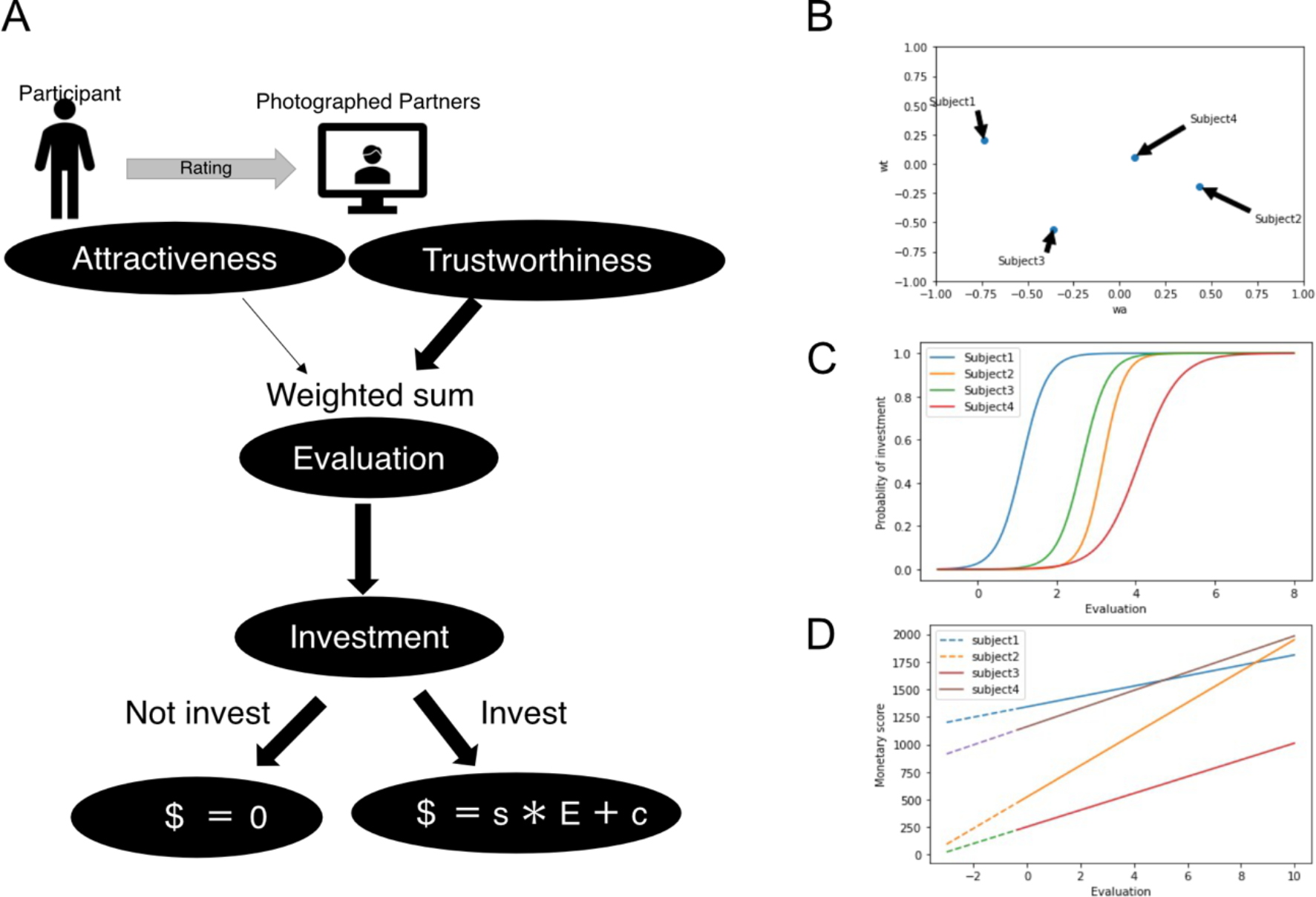
Decision-making model in trust game. **(A)** Decision-making process toward investment. The participants’ ratings for the photographed partners, investments, and monetary scores are observed, whereas the model parameters are unknown.**(B)** Weights for attributes of photographed partners are participant specific. **(C)** Decision of whether to make an investment. Participants tend to make an investment for highly evaluated photographed partners. The probability of investment obeys subject-specific sigmoid function. **(D)** Decision of the investment amount. If the participant decides to invest, his/her monetary score to invest linearly increases with the evaluation of the photographed partners in a participant-specific manner. Otherwise, the monetary score is zero.

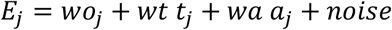

where *a*_*j*_ and *t*_*j*_ indicate the ratings of trustworthiness and attractiveness of the *j*-th photographed partner, respectively; *wo, wa*, and *wt* indicate the weights of the basal evaluation, attractiveness, and trustworthiness, respectively. The participant decides whether to invest, based on the following probability:

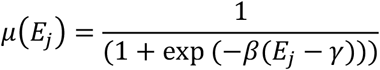

where *β* and *γ* indicate constant parameters, which control randomness and threshold of investment, respectively. If a decision to invest is made, the amount of money invested is determined as

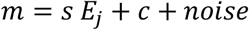

where *s* and *c* are constant parameters. If a decision to not invest is made, then the amount of money invested is zero. Furthermore, we assume that all parameters (i.e., *wo, wa*, and *wt*) are participant specific (**Fig. 3B-D**) and sampled from prior distributions (see Methods for details). Summarily, each participant in our model has their own weights for trustworthiness and attractiveness in the evaluation of partners’ faces. After evaluating their partner’s face, the participant decides whether to invest based on the weighted evaluation. If they decide to invest, a monetary score is determined through a weighted evaluation.

### Biased evaluation by patients with MMD

Before applying the Bayesian hierarchical model to the real data of the trust game, we noticed that some participants did not take the game seriously, with biased face evaluations and/or investments. Some participants always made the same evaluation for different partners’ faces (**Fig. 4A, B**) and invested the same amount of money in all partners. Conversely, other participants exhibited a more diverse range of choices (Fig. 4 B, C). In traditional psychological surveys, biased answers such as choosing only one or few options out of many according to subjective criteria are considered invalid. Thus, we quantified such biased answers for each participant using Shannon entropy ^16^, which is used in information theory to measure the variability of variables; high and low Shannon entropies represent varied and biased answers, respectively.

**Fig. 4:**
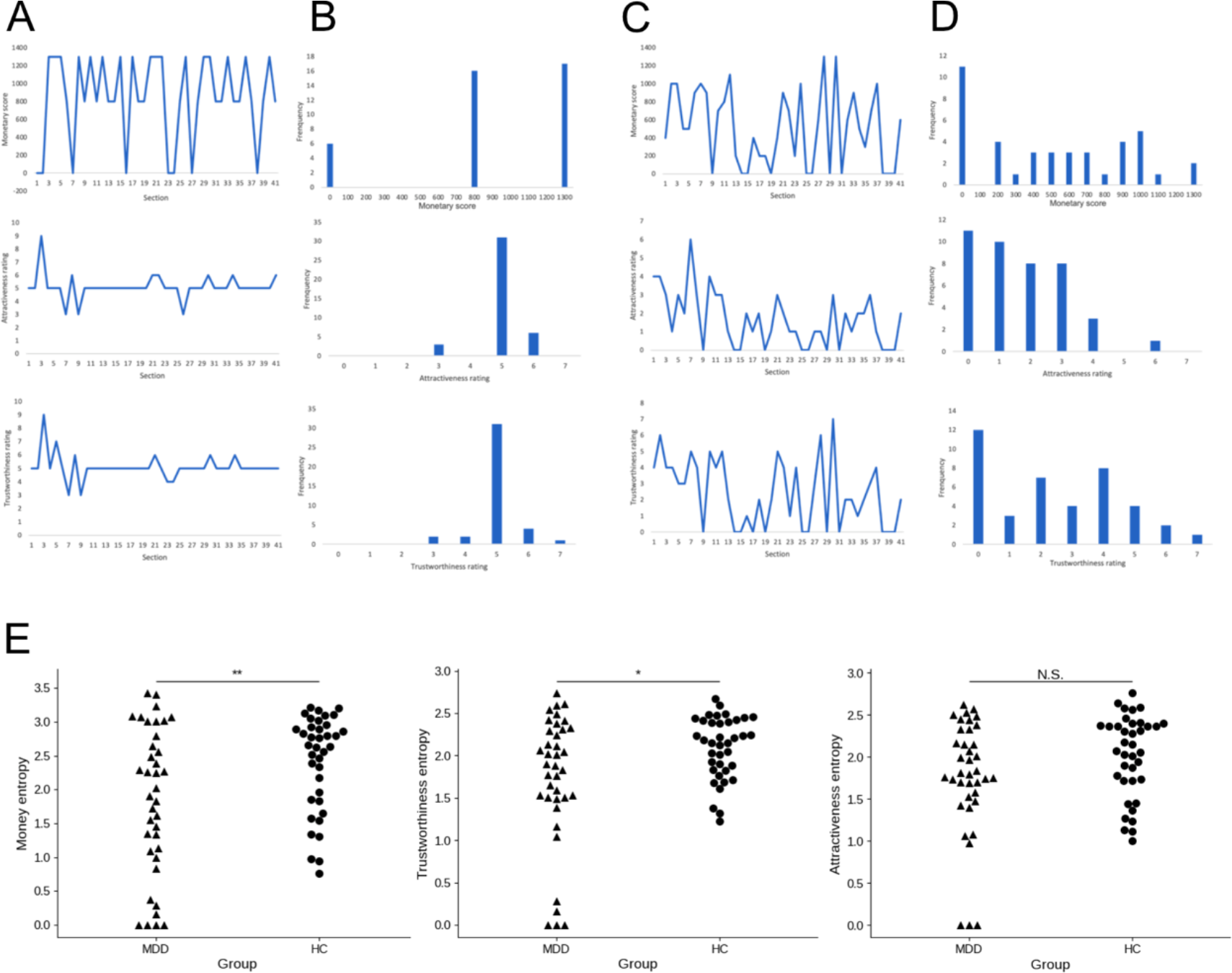
Biased, non-serious behaviors in trusting game. **(A)** Representative participants showing non-serious responses of attractiveness/trustworthiness ratings and monetary score of investment. **(B)** Histogram of each response for the representative participants in (A). **(C)** Representative participants showing diverse range of choices of attractiveness/trustworthiness ratings and monetary score of investment. **(D)** Histogram of each response for the representative participants in (C). **(E)** Comparison of Shannon entropy for each response between HCs and patients with MDD. Seriousness of participants for each response was quantified using Shannon entropy *S*. *p<0.05; **p<0.01; N.S. No significance. (Welch’s two-sample t-test).

We found that patients with MDD exhibited lower entropies of investment amount (p<0.01, **Fig. 4E**) and trustworthiness score (p<0.05) than HCs (**Fig. 4E**), whereas there was no significant difference in attractiveness between the two groups (**Fig. 4E**). The significant decrease in trustworthiness in patients with MDD suggests two possibilities: patients with MDD might be more likely to choose the same option without thinking about it, and they might not be sensitive to changes in facial information. Through information theory analysis, we identified answers with entropy (i.e., trustworthiness, attractiveness, investment) lesser than 1.5 as invalid answers, which were eliminated for the parameter estimation below.

### Estimation of participant-specific parameters

Based on the developed model, we aimed to determine the participant-specific parameters from the trusting behaviors (i.e., the participants’ scores of trustworthiness and attractiveness of their partners and investment amount) in the trust game. Using the MCMC (Markov chain Monte Carlo) method, we estimated all parameters from the training data (see Methods). We then demonstrated that the participant-specific models were able to predict whether and to what extent HCs and patients with MDD invested in their partners, which was confirmed by the test data (**Fig. 5A and B**).

**Fig. 5:**
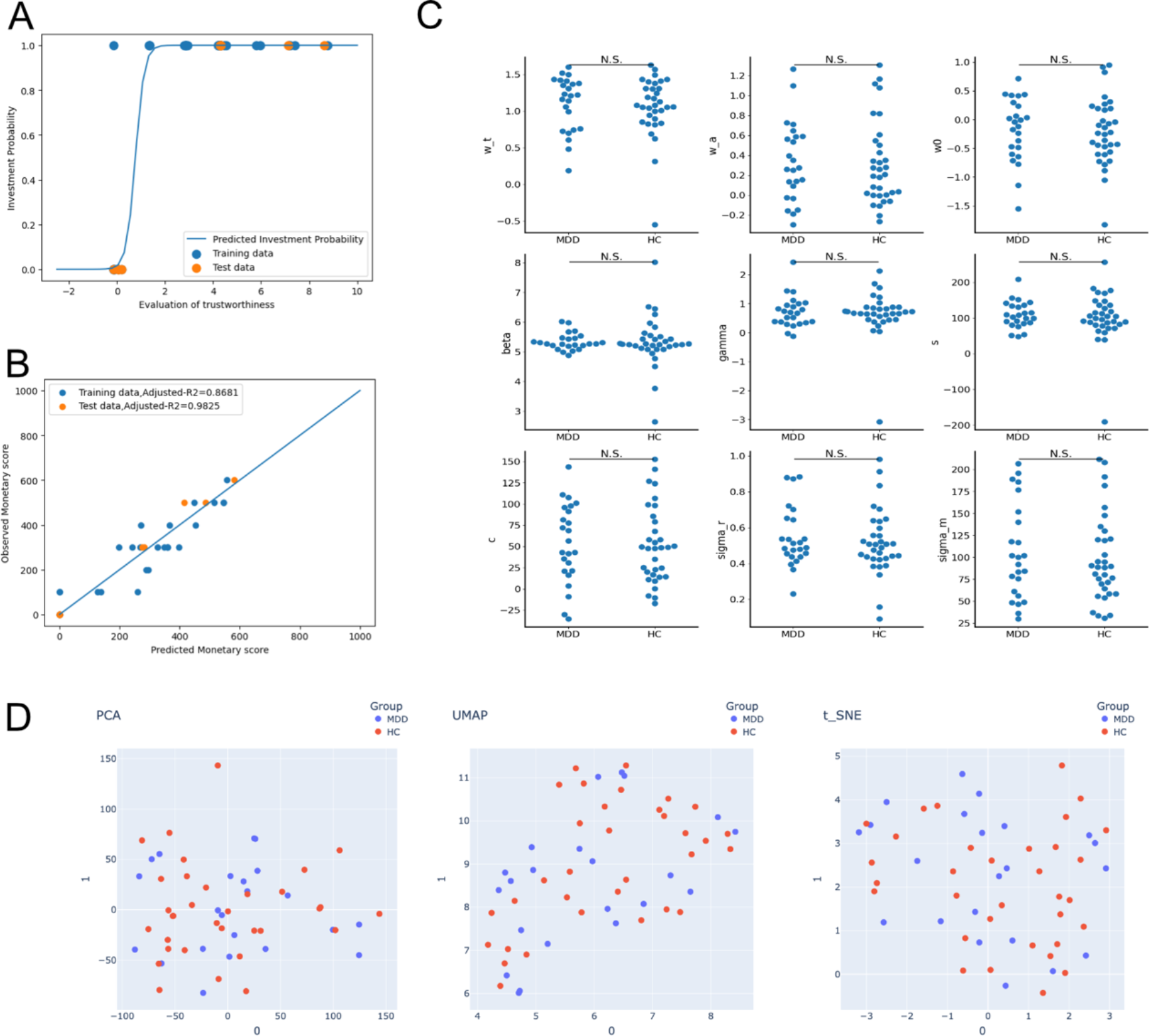
Prediction of participants’ behaviors in the trust game and parameter comparison between HCs and patients with MDD. **(A)** Predictions of whether each participant invests in the photographed partner. **(B)** Predictions of how much money each participant invests. **(C)** Comparison of the estimated parameters between HCs and patients with MDD. N.S. No significance. (Welch’s two-sample t-test). **(D)** Visualization of participants in model parameter space in 2D space. PCA, t-SNE, and UMAP were performed for dimension reduction of the estimated parameters.

We then examined how trust behaviors differed between the HC and MDD groups. Here, we used parameters estimated from the data for all 41 partners, which should be more reliable than those estimated from the partial training data. We found no significant differences in any of the nine parameters between the groups (**Fig. 5C**). In addition, we performed dimension reduction analyses (e.g., PCA, T-SNE, and UMAP) on a set of nine parameters, which characterized participant-specific trust behaviors (**Fig. 5D**). Contrary to our expectations, there were no clear boundaries between the parameters of the HC and MDD groups. These results suggest that participant-specific parameters are indicative of an individual’s personality and not related to the presence or absence of MDD.

### Uncovering Key Biomarkers Associated with Decision-Making in Trusting Behaviors

To examine whether <u>decision-making characteristics</u> were reflected by the metabolic state of each participant, we conducted a correlation analysis between participant-specific parameters and biomarkers (**Fig. 6A**). By summarizing the correlations (**Fig. 6A**), we noticed some biomarkers correlated with participant-specific parameters estimated from trusting behaviors (q value=0.05, false discovery rate), although the most of them were not correlated with trusting behaviors (**Fig. 2A**). We then identified 5ALA, acetyl carnitine, 2-aminobutyric acid, and glutamic acid, which differed from the identified biomarkers that correlated with the participants’ answers and behaviors. These four biomarkers negatively correlated with *wa*, a weighting parameter for attractiveness (**Fig. 6B**). 2-aminobutyric acid has been reported as a biomarker of depression ^17^. Acetylcarnitine is related to oxidative stress ^18^, and several studies have reported both positive and negative correlations with depression ^19,20^. Interestingly, acetyl carnitine is involved in the same metabolic pathway as acylcarnitine in the mitochondria, which we previously reported to be related to hikikomori ^11^. There are no reports on the function of 5ALA in neurons and depression. Glutamic acid, known as an excitatory neurotransmitter, is converted into gamma-aminobutyric acid (GABA), and abnormalities in the GABA neurotransmitter system have been observed in patients with mood and anxiety disorders. Thus, glutamic acid may be associated with MDD^21^.

**Fig. 6:**
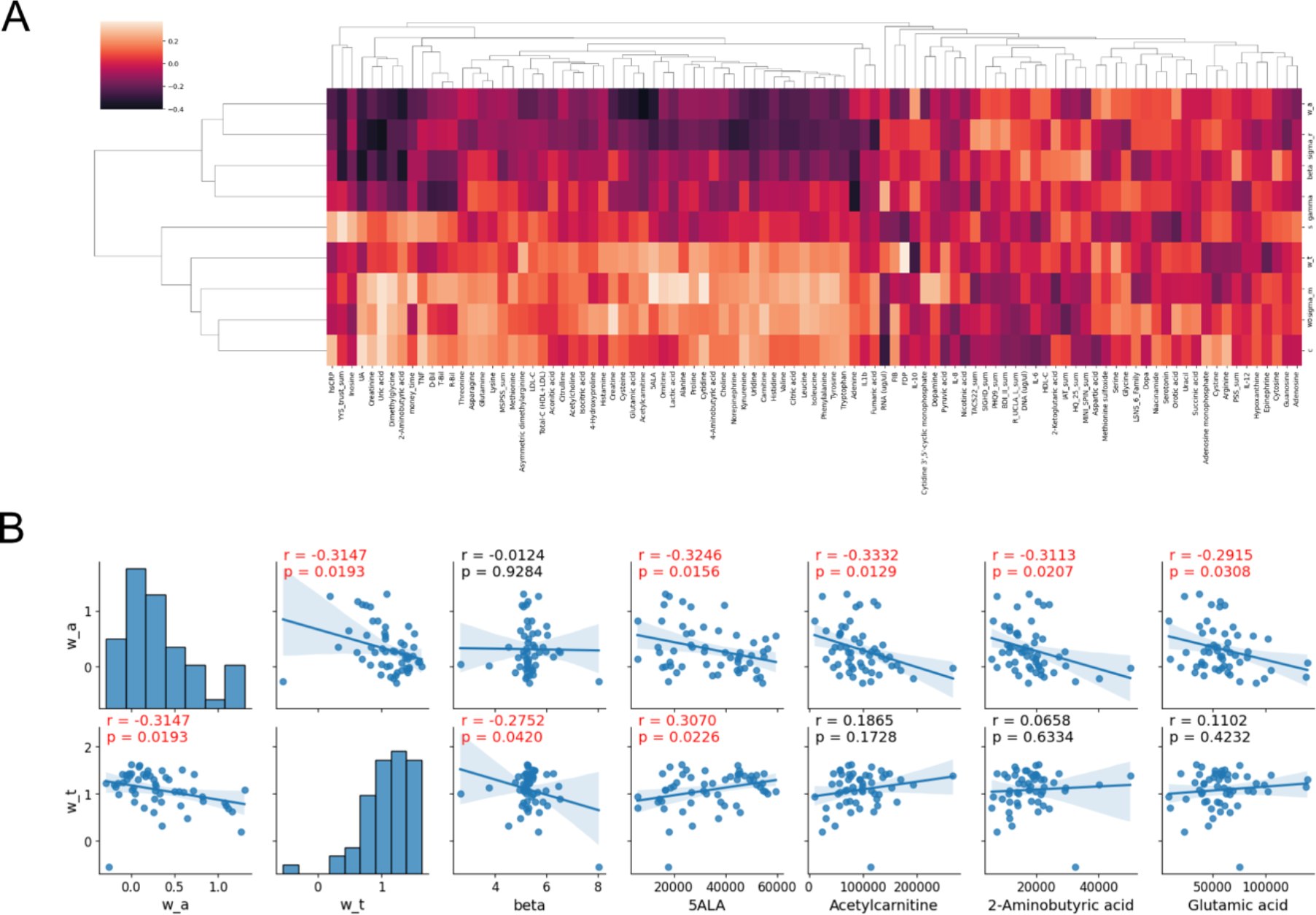
Screening of blood biomarkers correlated with the decision-making properties. **(A)** Heatmap of Pearson correlation coefficients between all estimated parameters and all measured blood metabolites. Hierarchical clustering was performed. Solid lines and shaded regions represent the estimated means and their 95% confidence intervals based on simple regression analysis, respectively.**(B)** Scatter plot of parameters and biomarkers for selected pairs. Correlation coefficients with absolute values > 0.3 and FDR < 0.05 were considered significant.

## Discussion

In this study, we performed a trusting game to obtain behavioral data on patients with MDD and healthy controls and measured metabolites from their blood samples. To simulate the trusting behavior of each participant, we developed a Bayesian hierarchical model. By estimating the model parameters for each participant, we identified several metabolites as biomarkers that correlated with the estimated parameters of the decision-making model. Our findings suggest that our approach can be of great help in understanding decision-making regarding trusting behaviors of individuals, including patients with MDD, and can provide new insights that explain the biological basis of trust behavior.

### Trusting behaviors in psychiatric disorders and hikikomori

We hypothesized that patients with psychiatric disorders might have a weak level of trust in their surroundings and adopted a trust game to assess their trust-based behaviors. The psychiatric patients partially co-exhibit symptoms of “withdrawal,” a lack of contact with the outside world. Therefore, modeling trusting behaviors is important to evaluate the decision-making process quantitatively.

In this study, some participants repeated the same answers and investments (**Fig. 4**). Such biased outcomes reflect the fact that the participants refused to think and provide valid answers. By evaluating the biases based on Shannon’s information entropy (**Fig. 4**), we found that patients with MDD had biased answers and investments compared with HCs, which is consistent with reports that patients with MDD exhibit autistic behavior^22^. Thus, participants with such biases provided less information for further analysis of their trusting behaviors.

### Biological function of the identified biomarkers

In this study, we proposed three levels of biomarkers: those discriminating psychiatric states, those predicting participants’ behaviors (i.e., answers and investments), and those associated with decision-making processes.

Previously, we identified several biomarkers for discriminating between patients with MDD and HCs at the first biomarker level. For example, increased levels of C-reactive protein (CRP), interleukin-6 (IL-6), and tumor necrosis factor-alpha (TNF-α), and decreased levels of brain-derived neurotrophic factor (BDNF) and serotonin, have been found to differentiate between patients with MDD and HCs ^23–26^.

Herein, we identified several biomarkers that were highly correlated with behavior (i.e., answers and investment) as secondary biomarkers. For example, participants with higher concentrations of uric acid, carnitine, methionine, creatinine, and nicotinic acid scored higher on the trustworthiness and attractiveness of the photographed partners and invested more in their partners (**Fig. 2**). Although we did not measure oxytocin (OT) in blood, it is known to promote perceived facial trustworthiness and attractiveness ^27^. However, such biomarkers, including OT, do not reflect how the subject made decisions based on external inputs, that is, the transformation from the facial evaluation of partners to investment.

At the third level of biomarkers, we identified several biomarkers in the blood associated with DMP. Acetylcarnitine is associated with the extent of attractiveness and oxidative stress. However, as it is well known, the concentration of metabolites in blood varies dramatically and can be influenced by numerous objective factors, such as sleep and diet. Therefore, further research is needed to determine how changes in acetylcarnitine concentrations affect changes in brain activity.

### Comparison with previous studies

Previous studies using fMRI have examined the neural basis of trusting behaviors and shown that trusting behaviors are associated with the activities of specific brain areas, such as the medial prefrontal cortex (mPFC) and temporo-parietal junction (TPJ) ^28,29^. In social implementations, it is impractical to use expensive fMRI to assess trust behavior. Our study aimed to use blood biomarkers to predict decision-making traits and trusting behaviors of participants. The biomarkers identified in this study enabled us to characterize DMP in a cost-effective manner.

There are several mathematical modeling studies on decision making in the trust game, compared to our model, which has the following advantages ^30–34^: First, unlike models in previous studies, our model considered the subjective perception of a partner’s photograph, that is, the attractiveness and trustworthiness of the partner. Generally, mathematical modeling in a trust game rarely focuses on the effect of subjective perceptions on decision making. On the other hand, in our daily life, our trust behaviors are largely influenced by the appearance of partners. Therefore, our mathematical model can assess the reality of trust decisions. Second, we solved the inverse problem, that is, the estimation of the model parameters from the behavioral data for each participant in a personalized manner. All previous studies developed forward models that needed to be simulated using manually assigned parameters. Thus, this study had the advantage of quantitatively assessing personality traits. Third, our modeling approach relates trusting decisions to biological mechanisms.Many studies on biomarkers and mathematical models have been conducted independently; however, our study integrated them. As the model parameters can be intuitively interpreted as decision-making properties, we can examine which blood metabolites correlate with the decision-making properties.

### Future perspective

In the future, it will be possible to use deep generative models to generate facial photos of partners by quantitatively manipulating their trustworthiness and attractiveness. With recent developments in deep learning, Generative Adversarial Network (GAN) has been used to generate realistic facial images with quantitative attributes^35^. With this technique, we can suppress bias caused by subjective judgments (trustworthiness and attractiveness), which are included in our model. In addition, our model can be extended to a continuous series of trust games, whereas it addresses only one-shot trust games. For patients with MDD, how the outcome of the previous round (e.g., betrayed by a partner) affects investment behavior in the next round and how this effect differs from HCs are topics that will be studied in the future.

## Methods

This case-control study was approved by the Ethics Committee of Kyushu University (30-452) and was conducted in accordance with the Declaration of Helsinki. The original data used in the present study were previously reported ^36^.

### Participants

We enrolled drug-free patients with MDD and HCs (age- and sex-matched controls) in the present study. All participants were Japanese (Asian) and provided written informed consent prior to the study. Patients were recruited from Kyushu University Hospital and its affiliations (mainly outpatient clinics). HCs were recruited via flyers at Kyushu University Hospital and the university campus. Structured Clinical Interview for DSM-IV-TR (SCID) was conducted by trained psychiatrists to diagnose MDD. We selected 38 drug-free patients with MDD (22 men and 16 women). As exclusion criteria for patients with MDD, we confirmed that none of the patients had a history of neurodegenerative diseases, psychotic disorders, mental retardation, substance abuse, or physical diseases, such as cardiovascular diseases, liver and kidney diseases, infectious diseases, malignant tumors, or head trauma. We selected 38 healthy participants as HCs through interviews based on the SCID regarding any previous or ongoing psychiatric disorders, physical diseases, and medications, which were set as the exclusion criteria.

### PC-based Trust Game

As introduced in our previous reports^3,12^, we developed an original PC-based trust game, which was conducted using a laptop to evaluate participants’ trusting behaviors and preferences for others. The participants (trustors) were instructed on the game rules. Subsequently, they were required to make decisions regarding the amount of 1,300 JPY (about 12 USD) to give to each of the 40 partners (trustees) and to rate the partners’ attractiveness based on facial photographs presented on a computer screen (score ranges: 0-9), which is thought to reflect the raters’ subjective preference for others (Preference Score). The amount of money given to the partner by the participant (Monetary Score) is tripled, and the partner then decides whether to split the money equally with the participant or take the entire amount. The participant’s decision on the amount of money to give to the partner is thought to reflect their level of trust in the partner.

In the experiment, the partners were virtual players on a computer screen. The participants had no information about their partners except for facial photographs, including the head and shoulders,with a neutral facial expression. Photos of the partners were selected from professional fashion models (i.e., “high-attractive partners”) or lay individuals (i.e., “ordinary-attractive partners”). We randomly selected 40 pictures (10 each of professional male fashion models, professional female fashion models, lay males, and lay females) for the trust game. Hence, the faces of the photographed partners had different levels of attractiveness. Participants were unaware of their partners’ decisions. After the experiments, each participant was paid an amount of money corresponding to the result of a randomly selected game as a reward.

### Blood biomarkers

All peripheral venous blood samples were collected between 10:00 and 15:00. Plasma was immediately extracted, frozen, and stored at -80°C until analysis. The measured blood biomarkers included routine blood biochemical markers and metabolites (metabolomics).

Routine blood biochemical markers including serum total-cholesterol (Total-C), high density lipoprotein-cholesterol (HDL-C), low density lipoprotein-cholesterol (LDL-C), fibrinogen (Fib), fibrin/fibrinogen degradation products (FDP), total-bilirubin (T-bil), direct-bilirubin (D-bil), indirect-bilirubin (I-bil), uric acid (UA), and high-sensitivity C-reactive protein (hsCRP) were measured by automatic biochemical analyzer (SRL, Inc., Tokyo, Japan). Plasma metabolites were measured by liquid chromatography-mass spectrometry (LC-MS) using LCMS-8060 (Shimadzu Corp., Kyoto, Japan), as previously described ^37^.

### Generative model for trusting behaviors

Our Bayesian hierarchical model assumes that the evaluation of the *j-*th photographed partner by the *i*-th subject, *E*_*i,j*_, is determined by the weighting of trustworthiness and attractiveness as follows:

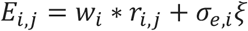

where *r*_*i,j*_ = (*a*_*i,j*_, *t*_*i,j*_, 1)^*T*^, *a*_*i,j*_, and *t*_*i,j*_ indicate the i-th participant’s rating of trustworthiness and attractiveness for the *j*-th photographed partners, respectively, and 1 represents common input among all partners; *w*_*i*_ = (*wa*_*i*_, *wt*_*i*_, *wo*_*i*_)^*T*^ indicates the weight vector for attractiveness, trustworthiness, and common input; *ξ* and *σ*_*e,i*_ indicate Gauss noise with zero mean and unit variance and its noise strength, respectively. Thus, *E*_*i,j*_ obeys the Gaussian distribution

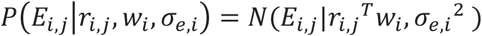

where *N*(*x*|*μ, σ*^2^)represents a Gaussian distribution with mean *μ* and variance *σ*^2^. The participants probabilistically decide whether to invest in a Bernoulli distribution.

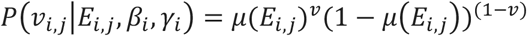

where *v* indicates binary variables; that is, 1 and 0 correspond to investment and no investment, respectively. *μ*(*E*_*i,j*_) is sigmoidal function:

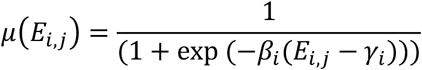

where *β*_*i*_ and *γ*_*i*_ indicate participant-specific constants that control for randomness and threshold of investment, respectively. If the investment is decided, the participant decides the amount of money as:

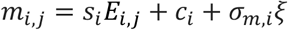

where s_*i*_ and *c*_*i*_ are participant-specific constants. If the participant does not select an investment, the amount of money is zero; thus, *m*_*i,j*_ obeys the following distribution:

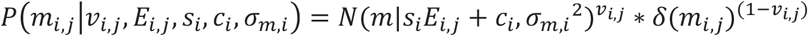

where *δ*(x) indicates the Dirac delta distribution.

### Prior distributions

For parameter estimation in a Bayesian manner, we introduced the following prior distributions:

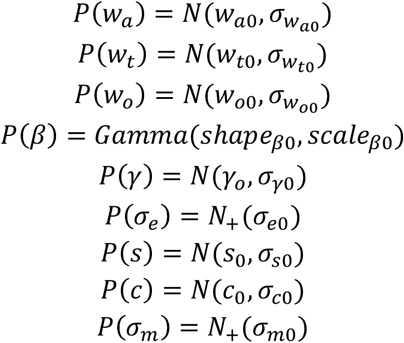

where *N*(*x*|*μ, σ*^2^) denotes a normal distribution with mu as the mean and sigma as the variance, *Gamma*(*a, b*) indicates a Gamma distribution with shape parameter *a* and rate parameter *b, and N*_+_(σ) indicates a half-normal distribution with scale σ. For most parameters, we used a normal distribution as the prior distribution. Because of the need to restrict beta to positive values, we chose the gamma distribution as the prior distribution. For the prior distribution of the variance parameter, we used a commonly used half-normal distribution of the variance parameter.

In the MCMC, we used *w*_*a*0_ = *w*_*t*0_ = *w*_*o*0_ = 0, 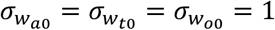, *shape*_*β*0_ = 5,*scale*_*β*0_ = 1, *γ*_0_ = *σ*_*γ*0_ = 1, *σ*_*e*0_ = 0.5, *s*_0_ = *σ*_*s*0_ = *c*_0_ = *σ*_*c*0_ = 100, and *σ*_*m*0_ = 100.

### Bayesian inference of parameters

We computed the posterior distribution of the parameters as

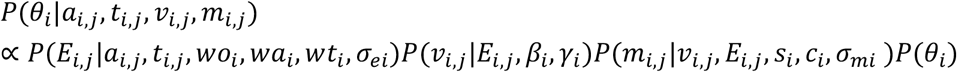

where *θ* = {*wa, wt*, … } and *P*(*θ*_*i*_) indicate prior distribution as *P*(*wo*_*i*_)*P*(*wa*_*i*_)*P*(*wt*_*i*_)*P*(*σ*_*ei*_)*P*(*β*_*i*_)*P*(*γ*_*i*_)*P*(s_*i*_)*P*(*c*_*i*_)*P*(*σ*_*mi*_). The inference of this distribution cannot be computed analytically; therefore, we employed Markov Chain Monte Carlo (MCMC) methods to perform the estimation. We used the No-U-turn sampler (NUTS) to sample this distribution, where 2,000 samples were drawn for burn-in and another 4,000 samples were drawn to estimate the distribution.

### Cross-validation

In the trust game, participants rated and invested in 41 partners. For each participant, we optimized the model using the ratings and investment amounts of 31 partners as the training set and obtained their specific parameters. In the participant-specific model, we predicted the investment and monetary scores of the remaining ten partners. We calculated the regression coefficient r^2 to estimate the credibility of the model.

## Acknowledgements

We are grateful to Prof. Michiyuki Matsuda for providing the research environment. This work was partially supported by a Grant-in-Aid for Scientific Research: (1) The Japan Agency for Medical Research and Development [AMED; JP21wm0425010 to T.H., T.A.K., and H.N.], (2) Cooperative Study Program of Exploratory Research Center on Life and Living Systems (ExCELLS) [program number 19-102 to H.N.], (3) The Japan Society for the Promotion of Science [KAKENHI; JP18H04042, JP19K21591, JP20H01773, and JP22H00494, 23H01044 to T.A.K.], and (4) The Japan Science and Technology Agency CREST [JPMJCR22N5 to T.A.K.]. The funders played no role in the study design, data collection and analysis, decision to publish, or manuscript preparation.

## Author Contributions

Z.C., H.N. T.H. and T.A.K. conceived the project. Z.C., Y.Y. and H.N developed the model. D.S., D.M.N., T.M., M.W. and T.A.K. conducted the experiments. Z.C. analyzed the data. Z.C. and H.N. wrote the manuscript with input from all the authors.

